# A fast and reliable larval sampling method for improving the monitoring of fruit flies in soft and stone fruits

**DOI:** 10.1101/2023.10.18.562869

**Authors:** Ghais Zriki, Rémy Belois, Christine Fournier, Léa Tergoat-Bertrand, Pierre-Yves Poupart, Amélie Bardel, Benjamin Gard, Nicolas O. Rode

## Abstract

The spotted-wing drosophila, *Drosophila suzukii* (Matsumura) (Diptera: Drosophilidae), threatens both the soft-skinned and stone fruit industry in Asia, Europe and America. Integrated pest management requires monitoring for infestation rates in real-time. Although baited traps are widely used for field monitoring, trap captures are weakly correlated to infestation rates. Thus, monitoring for larvae instead of adult flies represents the most reliable monitoring technique. Current methods for larval monitoring (e.g. dunk flotation) are both time-consuming and labor-intensive. In this study, we develop the sleeve method, a new method for monitoring larval infestation in strawberries through the inspection of fruits individually crushed within transparent plastic sleeves. Based on count data from both specialist and non-specialist observers, the estimation of larval infestation with the sleeve method is fast, precise and highly repeatable within and among all observers. Mean processing time is twice faster than previous methods and varies from 33 to 80 seconds per sample, depending on infestation levels. As the accuracy of the sleeve method decreases with infestation levels, we suggest ways to improve its accuracy by incubating fruits for 48h and calibrating data with exact counts. This new method is easier to use, faster and more cost-effective than previous monitoring methods. It could be used to monitor *D. suzukii* infestations in various fruit crops and is scalable to the farm-level for fruit growers or for academic research. Finally, the method represents a promising monitoring tool for effective management programs of *D. suzukii* and other insect pests of soft and stone fruits.

## Introduction

The Spotted Wing Drosophila (SWD), *Drosophila suzukii* (Matsumura) (Diptera: Drosophilidae), is a major insect pest of both soft and stone-fruits (e.g. strawberries, raspberries, blackberries, blueberries and cherries; Asplen et al. 2015). The serrated ovipositor of SWD females enables them to lay their eggs in healthy fruits, making them unmarketable. The rates of SWD infestation in soft fruits range from 20% to 40% and result in important revenue losses for farmers (Bolda et al. 2010, Cini et al. 2012).

SWD management relies mostly on chemical control using broad spectrum insecticides (Schetelig et al. 2018). Due to the toxicity of insecticides for human health and for the environment (Nicolopoulou-Stamati et al. 2016, Eggleton 2020), alternative biological control methods are currently under development (e.g. using natural enemies or the Sterile Insect Technique; Lee et al. 2019, Tait et al. 2021, Homem et al. 2022).

Integrated Pest Management (IPM) programs of SWD require an early and accurate detection of infestations to initiate chemical treatments (Tait et al. 2021). Baited and lured traps are widely used for SWD field monitoring (Cloonan et al. 2018). However, trap captures with baits are not or weakly correlated to infestation rates (Harris et al. 2014, Burrack et al. 2015). Thus, monitoring for larvae instead of adult flies represents the most reliable monitoring technique (Van Timmeren et al. 2017, 2021).

Few methods are available to estimate the prevalence and intensity of SWD infestation in soft- or stone fruits (Van Timmeren et al. 2017, 2021, Shaw et al. 2019, Babu et al. 2023). First, while the dissection of fruits allows the estimation of infestation prevalence, this method can underestimate infestation intensity, even when using a stereomicroscope (Shaw et al. 2019). Second, the extraction of larvae using fruit dunk floatation and filtration allows for the detection of small larvae resulting in more accurate estimation of both the prevalence and intensity of infestation than dissection methods (Van Timmeren et al. 2017, 2021). For experimental field trials that require large sample sizes, flotation and filtration can be labor intensive and require large quantities of liquid for floatation (i.e. either sugar, salt or detergent solution; Van Timmeren et al. 2017). Although its does not require squeezing fruits, an alternative method based on the vacuum extraction of larvae requires buying a vacuum pump (for approximately $250) and operating it safely (Babu et al 2023). As all these methods can be impractical for the daily inspection of fruit infestation by consultants or fruit growers, cheaper, more practical and easier to use methods are highly needed.

To estimate the prevalence and intensity of infestation by *D. suzukii*, we developed the sleeve method, an easy-to-use method based on fruit crushing in plastic sleeves and direct observation. We estimated the performance of the sleeve method using several criteria (failure to detect an infestation, false detection of an infestation, speed, accuracy, precision, within- and among-observer repeatability) using infested strawberries. Due to the fiber content of strawberry, the estimation of infestation rates in this fruit is more challenging than estimation in other fruits (blueberry, blackberry, raspberry, and cherry; Shaw et al. 2019). We show that the sleeve method is quicker and likely more precise than previous methods.

## Materials and Methods

### General methods

All experiments were conducted at Centre Technique Interprofessionnel des Fruits et Légumes (CTIFL) in Bellegarde, France in 2021-2022. Additional details regarding the sleeve method are provided in Supplementary Materials (Fig. S1).

#### Sample preparation

We first estimated the average time required to prepare samples using 99 commercial strawberry fruits. After removing the calyx, we placed each fruit in the middle of an A4 transparent plastic sleeve (Fig. 1A). We then softly smashed each fruit into a thin layer of puree with the hand palm without applying too much pressure (Fig. 1B).

**Fig. 1.**
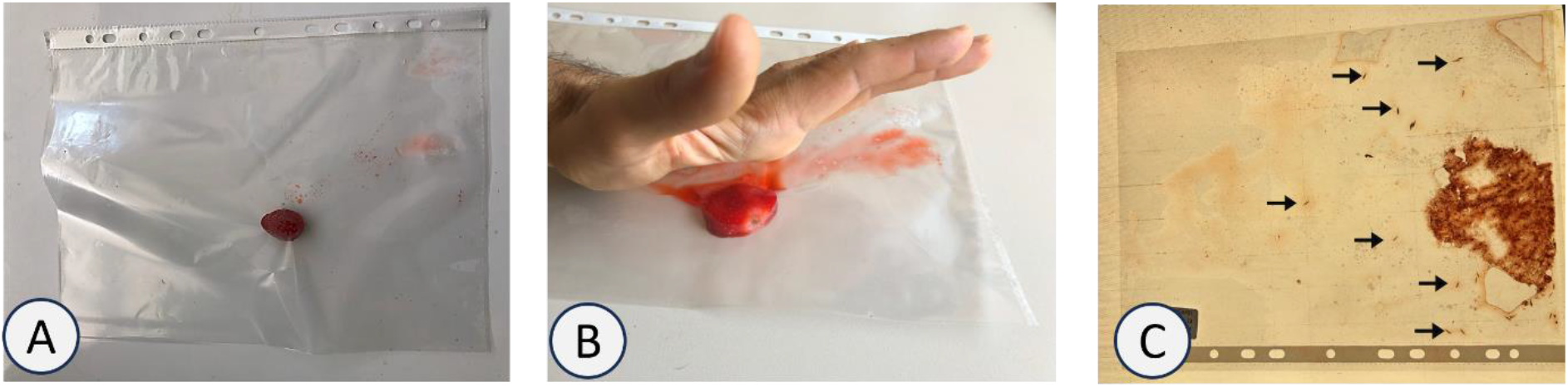
Monitoring the prevalence and intensity of infestation of *D. suzukii* larvae in strawberries using the plastic sleeve method. (A) place one or more strawberry in the middle of a transparent plastic sleeve, (B) smash the strawberry with hand palm, to count larvae (C) place the sleeve against a light source. Larvae are indicated by arrows.

#### Larval count

To estimate the performance of the sleeve method with fruits naturally infested with *D. suzukii*, we sampled 30 strawberries in a glasshouse. We tested whether incubating the samples increases the performance of the sleeve method, by incubating 15 strawberries at 24°C for 48h before freezing, while the other 15 strawberries were frozen directly without any incubation. After thawing the strawberries, we prepared each sample according to the abovementioned method. We inserted a transparent 1-cm grid paper between the plastic sleeve and a backlight source to facilitate counting (Fig. 1C). Two experienced observers (G.Z. and R.B.) estimated the exact number of larvae for each of the 30 samples (several larval counts of each sample). We then compared the performance of the method between 7 specialist and 3 non-specialist observers (with or without any previous experience with the observation of *D. suzukii* larvae) who counted the number of larvae in each of the 30 strawberry samples. Each observer counted the larvae in each sample twice (30 samples x 10 observers x 2 replicates = 600 observations in total). We recorded the time required for counting each count replicate.

We estimated the prevalence and intensity of infestation across the 30 strawberry samples. To estimate the rate of false negative and false positive of the sleeve method, we estimated the rate of failure to detect an infestation and the rate of false detection of infestation.

### Estimating the speed, accuracy, precision and repeatability of the sleeve method

To further assess the performance of the sleeve method, we used the dataset with 600 observations to estimate the speed, accuracy, precision and repeatability. We estimated the accuracy (bias relative to the exact number of larvae) and precision (magnitude of the standard deviation among observer) of each larval count. For each sample, and separately for specialist and non-specialist observers and for the first and second replicate counts, we computed the bias of each count as the difference between the mean of larval counts and the exact number of larvae. Similarly, we computed for each sample, separately for specialist and non-specialist observers and for the first and second replicate counts, the standard deviation among larval counts (2 types of observers x 2 replicates x 30 samples = 120 bias/standard deviation estimates in total).

Finally, we also estimated the within and among-observer repeatability of larval counts (Nakagawa and Schielzeth 2010). Indeed, the magnitude of the precision should be compared to the overall variance among samples. For example, let us imagine that a given sample is infested by 30 larvae and that two observers counted 29 and 31 larvae respectively. If the overall variance is large (e.g. with some fruits with no larva and some fruits with 100 larvae), the standard deviation of 1 larva among observers can be considered as irrelevant (high among-observer repeatability). In contrast, if the overall variance is small (e.g. with every fruits infested by either 29, 30 or 31 larvae), the standard deviation of 1 larva among observed makes the method useless (low among -observer repeatability).

### Statistical analyses

All statistical analyses were performed using R statistical software (R Core Team 2022). To estimate the speed, accuracy, precision and repeatability of the sleeve method, we used separate Linear Mixed Models (LMM) with a Gaussian distribution. For the analyses of speed, accuracy and precision, we tested for main effects and interactions using stepwise model selection with Likelihood Ratio Tests (LRT; Bates et al. 2014). Finally, for each analysis, we visually checked for the normality of the residuals of the final model.

To investigate the effects that could affect the speed of the sleeve method, we fitted LMMs on the log-transformed time to count each sample. Fixed effects included the 48 hour-incubation status (“yes” vs. “no”), the long-term experience of the observer (“specialist” vs. “non-specialist”), and the short-term experience of the observer (“first” vs. “second” replicate count) as factors and the expected number of larvae as a continuous covariate. To test whether the effect of incubation varied depending on the number of larvae, the model also included an interaction between incubation status and expected number of larvae. To account for the non-independence among observations from the same fruit sample or from the same observer, all models included the identity of the strawberry and the identity of the observer as random effects. To investigate the accuracy of the sleeve method, we fitted LMMs on the 120 bias estimates using the same four fixed effects as in the analysis above. To account for the non-independence among observations from the same fruit sample, all models included the identity of the strawberry as random effect (observations are averaged across observers so that a random observer effect is not necessary). To investigate the precision of the sleeve method, we fitted LMMs on the 120 standard deviation estimates using the same four fixed effects and the random effect as in the accuracy analysis above.

Finally, to estimate the within- and among-observer repeatability, we fitted a single LMM on the log-transformed count of larvae which included the identity of the strawberry sample and the identity of the observer as random effects (Eq. 11 in Nakagawa and Schielzeth 2010).

## Results

### Failure to detect an infestation and false detection of an infestation

The prevalence and intensity of infestation was 33% and 4.6 (SD=6.94) larvae per strawberry fruit in the experiment. Among the 20 samples with at least one larva, failure to detect an infestation ranged from 5% to 15% depending on the observer. Across observers, this rate was 11% (45 in 400 counts). Among the 10 samples with no larvae, only one observer counted one larva in one of the two replicate counts. Hence, the rate of false detection of infestation was 0.5% (1 in 200 counts).

### Speed of the sleeve method

The average time required to crush one strawberry in a plastic sleeve was 16 seconds (SD=11 seconds). The average time required to count larvae in one strawberry sample was 25 seconds (SD=15). Hence, the total time to process one strawberry sample was 41 seconds (SD=19). The average time required to count larvae in each fruit increased with the number of larvae (from 17.5 seconds for fruits with fewer than five larvae to 63.0 seconds for fruits with more than 15 larvae; 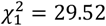 *P* = 5· 10^−8^; Fig. 2A; Table S1). When accounting for this effect of the number of larvae on count time, incubating fruits for 48 hours did not significantly decrease the amount of time required to count larvae in each fruit (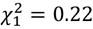 *P* = 0·64; Fig. 2A). Observers with previous experience in counting larvae did not count faster than observers without any previous counting experience (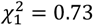 *P* = 0·39), indicating no long-term effect of previous experience. For each sample and observer, the time needed to count larvae was lower for the second than for the first count (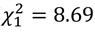 *P* = 0·003; Fig. 2A), indicating a short-term effect of previous experience.

**Fig. 2.**
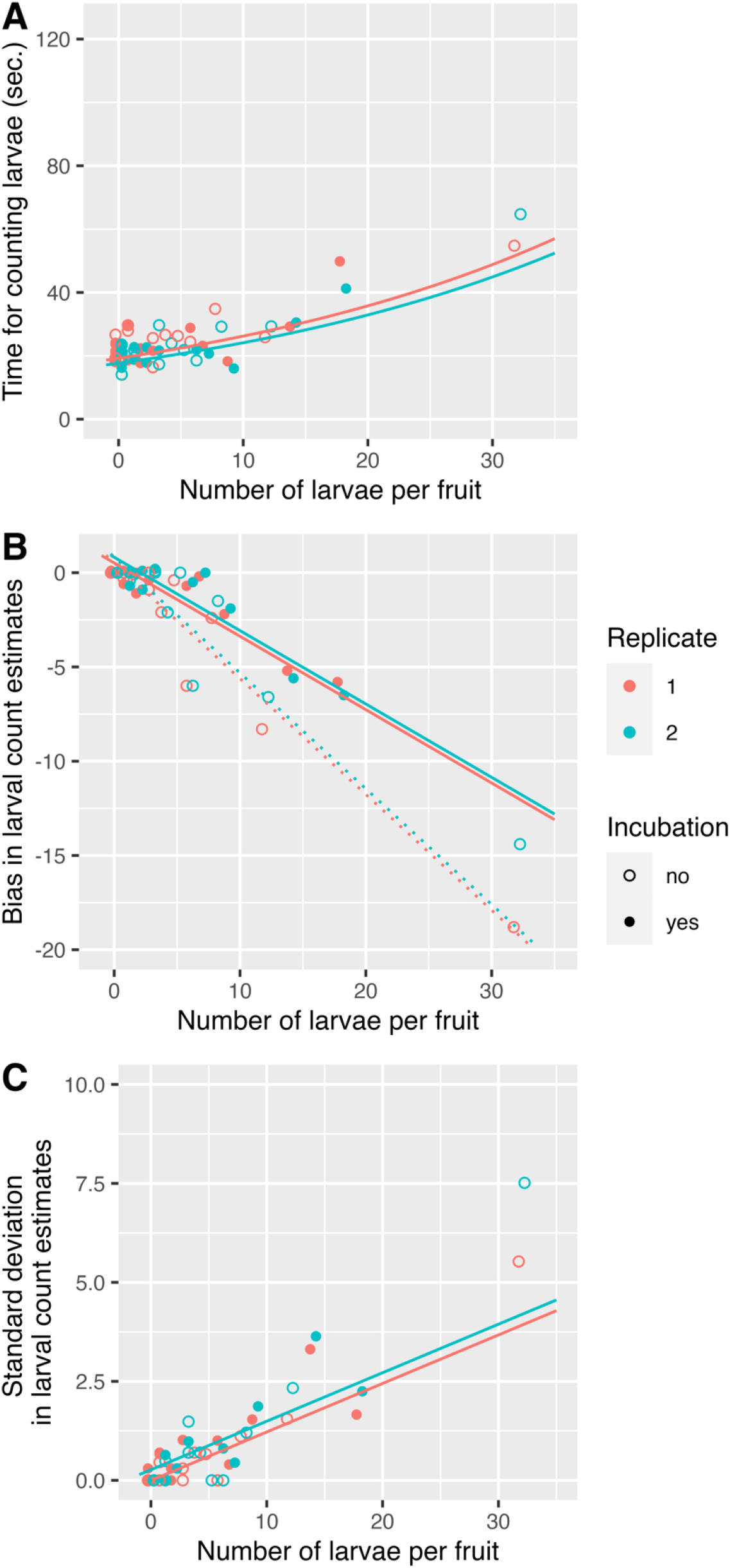
The three parameters for the performance of the sleeve method, (A) counting time, (B) accuracy and (C) precision (i.e. inverse of the standard deviation among estimates), can depend on the number of larvae, on the 48-hour incubation or not (filled and solid circles) and on the order of count replicates (red and green symbols). Dotted and solid lines represent the fitted lines for samples respectively counted with or without a 48-hour incubation (the two types of lines are superimposed in panels A and C).

### Accuracy of the sleeve method

The number of larvae per sample was consistently underestimated (negative bias) as the exact number of larvae per sample increased (Fig. 2B). The increase in this underestimation was significantly lower in incubated than in non-incubated samples (significant interaction between incubation status and exact number of larvae; 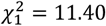 *P* = 0·0007). We estimated that when the exact number of larvae increased by 10 larvae, the observed number of larvae increased by only 5 larvae when samples are not incubated or by 3 larvae when samples are incubated (dotted and solid lines in Fig. 2B). Importantly, this interaction remained significant when removing one non incubated sample that included more than 30 larvae (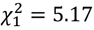 *P* = 0·02). For each sample and observer, the accuracy of second count replicate was slightly and significantly higher than that of the first replicate (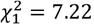 *P* = 0·007; red and green lines in Fig. 2B). Specialist observers did not count more accurately than non-specialist observers (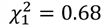 *P* = 0·41), indicating the absence of long-term effect of observer experience on the method’s accuracy.

### Precision, within- and among-observer repeatability of the sleeve method

The precision of the method decreases as the exact number of larvae per sample increases, as expected with count data (increase in the standard deviation with the number of larvae in Fig. 2C). Sample incubation did not significantly affect the precision of counts (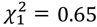 *P* = 0·42; superimposed solid and dotted lines in Fig. 2C). For each sample and observer, the precision of second count replicate was slightly and significantly lower than that of the first replicate (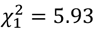 *P* = 0·01; red and green lines in Fig. 2C). However, specialist observers did not count more precisely than non-specialist observers 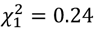 *P* = 0·62), indicating the absence of long-term effect of observer experience on the method’s precision.

The variance in the number of larvae among samples (V_*sample*_ = 0·81) was large relative to the variance among observed (V_*among*_ = 0·005) and the residual within-observer variance (V_*within*_ = 0·04). Hence, within- and among-observer repeatabilities were greater than 94% (Fig. S1).

## Discussion

We developed a new easy-to-use method for monitoring *D. suzukii* infestation in strawberries. By crushing fruits within a transparent plastic sleeve, the method provided satisfactory results for all performance criteria (failure to detect an infestation, false detection of an infestation, speed, precision, within- and among-observer repeatability), but accuracy at high infestation levels (Fig. 2B). Overall, the sleeve method allows for the fast detection of larvae with the naked eye or with a magnifier. The thorough experimental design and robust statistical framework used to assess the performance of the sleeve method is readily transferable to assess the performance of similar methods to count the larvae of other insect pests.

### Advantages of the sleeve method compared to other methods

The underestimation of the number of larvae at high infestation levels is likely due to observers losing track of the larvae they already counted and might be independent of the fruit preparation method. If intensities of infestation are to be estimated, we recommend counting the exact number of larvae on a fraction of the samples to perform a calibration. As a rule of thumb based on calibration from Fig. 2B, we estimate that the observed number of larvae should be multiplied by 1.5 (respectively by 2) to estimate the exact number of larvae in the presence (respectively in the absence) of incubation.

Compared to previous methods, the sleeve method allows higher recovery rates and is both faster and easier to use, everything else being equal (i.e. accuracy, precision, within- and among-observer repeatability of counts; Table 1). With the method, accuracy is not affected by larval recovery, as 100% of the larvae end up in the sleeve. Total processing time per fruit (to prepare samples and count larvae) is 41 seconds for infestation intensities ranging from 5 to 10 larvae, which results in a high throughput of ∼100 fruits per hour. Furthermore, the sleeve method requires less consumables than other methods, as transparent plastic sleeves are reusable many times (Table 1). The method is also versatile as unhatched eggs and larvae can be counted with the naked eye or with a stereomicroscope (the latter allows for a higher detection rate of eggs and small L1 or L2 larvae).

**Table 1.** Comparison of the different methods available to estimate the prevalence and intensity of fruit infestation by *Drosophila suzukii*. We assume that count error is low and equal among the different methods.

| Reference | Method | Mean/variance in prevalence and infestation intensity | Larval recovery | Ease of use | High-throughput | Stereomicroscope | Material required | Potential freezing for later detection |
| --- | --- | --- | --- | --- | --- | --- | --- | --- |
| Shaw et al 2019 | Dissection | Mean & variance | 100% | - | - | No/Yes | Dissection forceps | Yes |
| Shaw et al 2019 | Crush+ Floatation | Mean & variance | ≤100% | - | - | No | Plastic funnel, cloth, floatation solution | No |
| Van Timmeren et al 2017; Van Timmeren et al 2021 | Crush+ Floatation+ Filtering | Mean & variance | ≤100% | + | + | No/Yes | Plastic funnel, cloth, reusable coffee filter, floatation solution | No |
| Babu et al 2023 | Vacuum+ Rinsing+ Filtering | Mean only | ≤83% | + | ++ | No/Yes | Vacuum pump, reusable coffee filter | No |
| This study | (Incubate+)<br>(Freeze+)<br>Crush | Mean & variance | 100% | ++ | ++ | No/Yes | Plastic sleeve (reusable if properly rinsed with water) | Yes |

Alternative methods based either on fruit dunk floatation and filtration or vacuum extraction are characterized by larval recoveries lower than 100% as some larvae can remain within the fruit (Babu et al. 2023). This results in the underestimation of the intensity of infestation and a lower accuracy than with the sleeve method. For alternative methods, total processing time per fruit is typically 120 seconds or higher (Van Timmeren et al. 2021), which likely results in a lower throughput than the sleeve method. Finally, other methods require a liquid either for dunk floatation (Van Timmeren et al. 2021) or for rinsing fruits to collect the remaining larvae (Babu et al. 2023), which cannot be reused.

### Advantages of the short-term experience of observers

Although not investigated in other methods, previous short-term experience of observers allows faster and more accurate counts. Indeed, the accuracy of the sleeve method was on average higher in the second than in the first replicate count (Fig. 2B). This suggests that the skills of observers to count larvae could be improved by a quick preliminary session (e.g. by counting mock samples). Importantly, this session should occur just before counting the samples of interest, as the long-term experience of the observer had no effect on the speed, accuracy or precision of larval counts.

### Advantages and drawbacks of incubation in the sleeve method

In the sleeve method, incubating strawberries at 24°C for 48 hours allows for more accurate estimations of the intensity of fruit infestation, but not faster or more precise counts. Incubation likely allows the detection of larvae initially present as eggs or L1. Incubation is particularly advantageous for estimating the number of larvae per fruit when infestation intensities are high, as it reduces the underestimation of the number of larvae (Fig. 2B). Everything else being equal, incubation does not affect larval recovery, as 3rd instars within fruits are counted as pupae after 48 hours of incubation at ∼20°C. The main drawback of incubation is a potential underestimation of prevalence and intensity of fruit infestation due to larval mortality. For example, microbial development during incubation could decrease larval survival when using incubation in the sleeve method. The relative drawbacks of incubation due to larval mortality remain to be quantitatively assessed.

### Advantage for fruit growers and academic researchers

Previous methods have not been widely adopted by growers as they are time consuming, labor-intensive and require specific consumables (Tait et al. 2021). Fruit growers, academic researchers and consultants can easily use the sleeve method, as no specific training is required. Fruit growers are often interested in assessing infestation rates in real time and are not interested in infestation intensity. In this case, the incubation step of the sleeve method can be omitted.

In contrast, academic researchers are often interested in assessing both the mean and variance of the prevalence and intensity of fruit infestation (Table 1; McIntosh et al. 2022). In this case, incubation can allow for a more accurate estimation of infestation intensity. In addition, for experiments with large sample sizes, samples can be stored frozen until further processing. The sleeve method will be particularly useful for the development of innovative pest control strategies by allowing the accurate estimation of their efficiency in both laboratory and field settings (e.g. for the development of the sterile insect technique).

Based on data from both specialist and non-specialist observers, the sleeve method is more time efficient than previous methods. Incubating fruits improves the accuracy of the method. This study represents a first proof of principle of the utility of this new method for the estimation of the prevalence and intensity of infestation of soft- and stone-fruits by *D. suzukii*. As strawberries are among the fruits where the detection of infestation is the most difficult (Shaw et al. 2019), the sleeve method should easily be transferable to other fruits. The thorough experimental design and robust statistical framework used here will allow for additional development of the sleeve method to detect *D. suzukii* larvae in other fruits or to detect the larvae of other insect pests that infest soft- or stone fruits. It thus represents a promising alternative to floating techniques currently used to detect the larvae of other pest insects such as tephritids (Bactrocera tryoni, Ceratitis capitata or Rhagoletis sp.; Yee 2014, Balagawi et al. 2022).

Finally, we hope that the ease of the sleeve method will encourage fruit growers to incorporate direct monitoring of fruit samples in their IPM program. The accurate detection of larvae and other insect pests will likely help them make informed decisions and improve the timing of control methods.

## Data availability

The data and R scripts to analyse the performance of the sleeve method are available at: https://github.com/nrode/Y2023.LarvCount

## Supporting information

The sleeve method (average counting time of SWD larvae)

## Acknowledgements

We are grateful to the volunteers for their participation in the study and to Frédéric Soula for providing strawberries for the experiment. We also thank Joël Escaich, Isaack Wanko and Antoine Escamez for their help. L.T.B. and P.P. were supported by funding from the Agence Nationale de la Recherche (ANR-21-ECOM-0002).

## Author contribution

Conceptualization: G.Z., N.O.R.

Experimental design and data acquisition: G.Z., R.B., C.F., L.TB., PY.P. and A.B., with inputs from N.O.R.

Statistical analyses and writing: G.Z. and N.O.R with inputs from B.G.

