## Supplementary material for "A fast and reliable larval sampling method for improving the monitoring of fruit flies in soft and stone fruits": The sleeve method (average counting time of SWD larvae)

### The sleeve method

**Step1: sample preparation.** Collect fruits and remove the calyx with entomological forceps or by pulling it off with fingers (note that this step could be done after incubation, but it increases the risks of removing some fruit parts with the calyx).

Eggs and young larvae are difficult to detect with the naked eye. We recommend incubating strawberries for 48h-72h at 24°C (Fig. S1A) to allow for the development of eggs and young larvae into bigger larvae. Please note that incubation duration should be adjusted to the temperature used. Longer incubation periods can be used for lower temperatures. Higher temperatures allow for shorter incubation periods, provided they do not increase larval mortality rates.

Freezing fruits for at least 24 hours can ease the next steps of crushing fruits and counting larvae (live larvae can escape plastic sleeves). If counting is delayed, freezing can be done over longer time periods.

**Step2: fruit crushing.** Allow samples to thaw for one or two hours before crushing. Place each fruit in the middle of an A4 transparent plastic sleeve (large fruits can be placed further away from the opening of the pocket to prevent leaking) (Fig. S1B). Softly smash each fruit into a thin layer of puree with the hand palm without applying too much pressure (Fig. S1C). At this step, it is possible to freeze the pocket for a later counting.

**Step3: larval count.** Place the pocket on a glass window (Fig. S1D) or with a backlight (Fig. S1E) or under a stereomicroscope (Fig. S1F). We recommend using a transparent 1-cm grid over the light source to facilitate the counting of larvae. Larvae are clearly visible with the naked eye, especially with backlight.

**Table S1.** Average count time (seconds) of *Drosophila suzukii* larvae per fruit and by incubation status and level of experience of the observer. The number of larvae did not differ between incubated and non-incubated samples.

| Infestation intensity | Incubation | Previous Experience | Mean count error | Mean Time (sec) | SD | SD | Number of fruits |
|---|---|---|---|---|---|---|---|
| 0-5 | no | no | 0.35 | 17.4 | 7.07 | 0.91 | 60 |
| 5-10 | no | no | 2.78 | 23.12 | 10.05 | 2.37 | 18 |
| 10-15 | no | no | 8 | 28.35 | 11.2 | 4.57 | 6 |
| +15 | no | no | 18.83 | 52.18 | 17.84 | 7.28 | 6 |
| 0-5 | no | yes | 0.36 | 23.77 | 12.06 | 1.02 | 140 |
| 5-10 | no | yes | 2.74 | 27.02 | 12.8 | 1.97 | 42 |
| 10-15 | no | yes | 7.21 | 27.26 | 14.41 | 3.85 | 14 |
| +15 | no | yes | 15.64 | 62.98 | 29.04 | 7.76 | 14 |
| 0-5 | yes | no | 0.32 | 17.49 | 5.4 | 0.7 | 60 |
| 5-10 | yes | no | 1.06 | 21.01 | 9.57 | 2.26 | 18 |
| 10-15 | yes | no | 6.5 | 26.82 | 18.07 | 7.38 | 6 |
| +15 | yes | no | 5.5 | 54.55 | 23.53 | 9.61 | 6 |
| 0-5 | yes | yes | 0.26 | 22.98 | 11.09 | 0.94 | 140 |
| 5-10 | yes | yes | 0.95 | 21.68 | 9.17 | 1.42 | 42 |
| 10-15 | yes | yes | 5.07 | 31.2 | 20.33 | 5.43 | 14 |
| +15 | yes | yes | 6.43 | 41.67 | 27.49 | 7.35 | 14 |

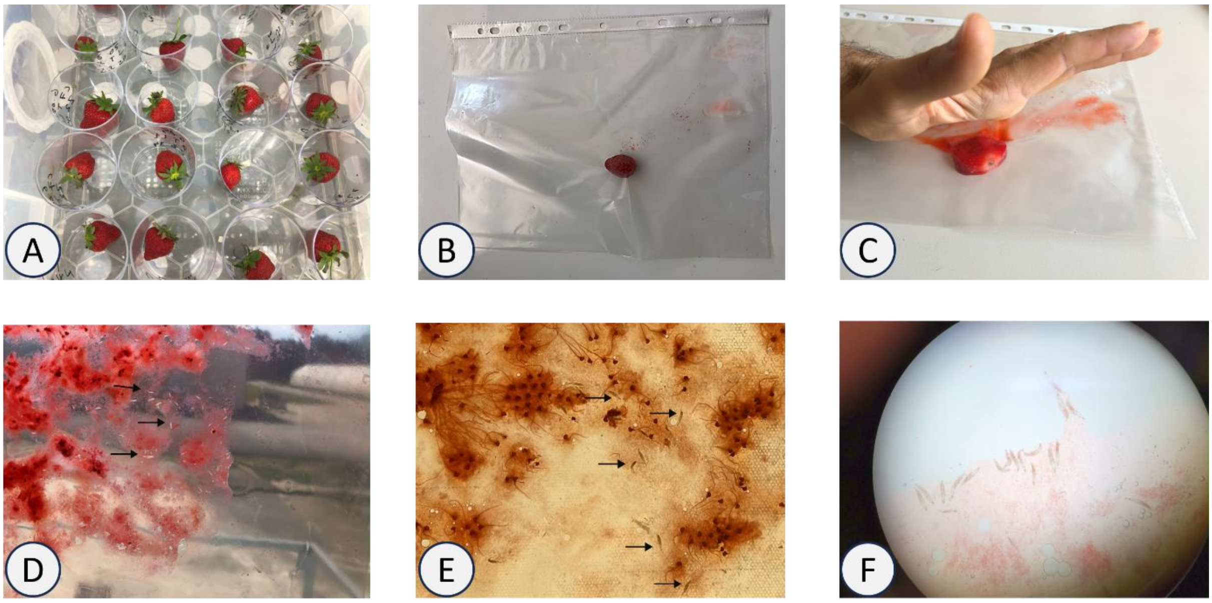


**Fig. S1.** Monitoring the prevalence and intensity of infestation of *D. suzukii* larvae in strawberries using the plastic sleeve method. (A) facultative preliminary incubation in plastic glasses at 24°C for 48 hours (enhance the accuracy of larval counts), (B) place one or more strawberry in the middle of a transparent plastic sleeve, (C) smash the strawberry with hand palm, to count larvae (D) place the sleeve against a window, or (E) against a light source or (F) under a stereomicroscope. Arrows indicate the larvae.

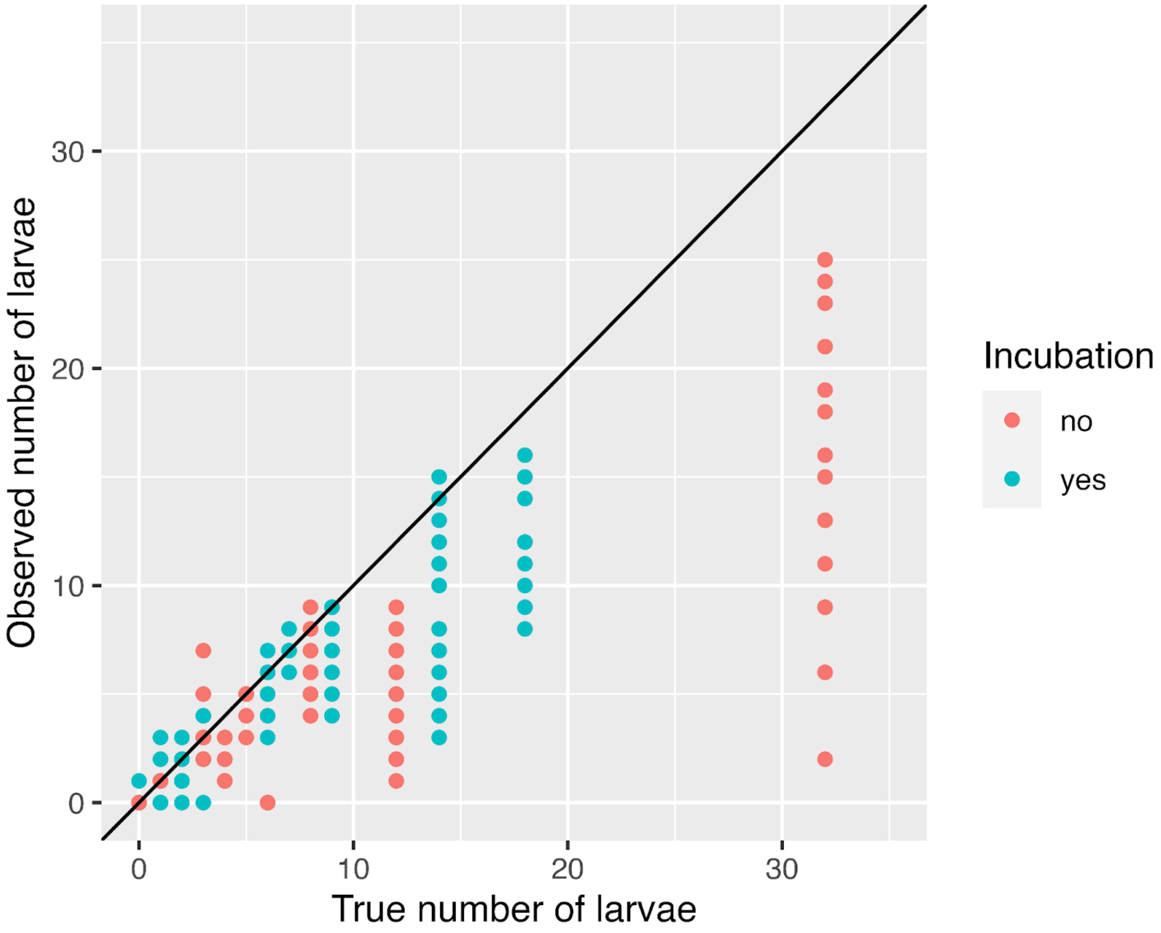


**Fig. S2.** Increase in bias and decrease in precision of larval count estimates with an increasing number of larvae within each fruit

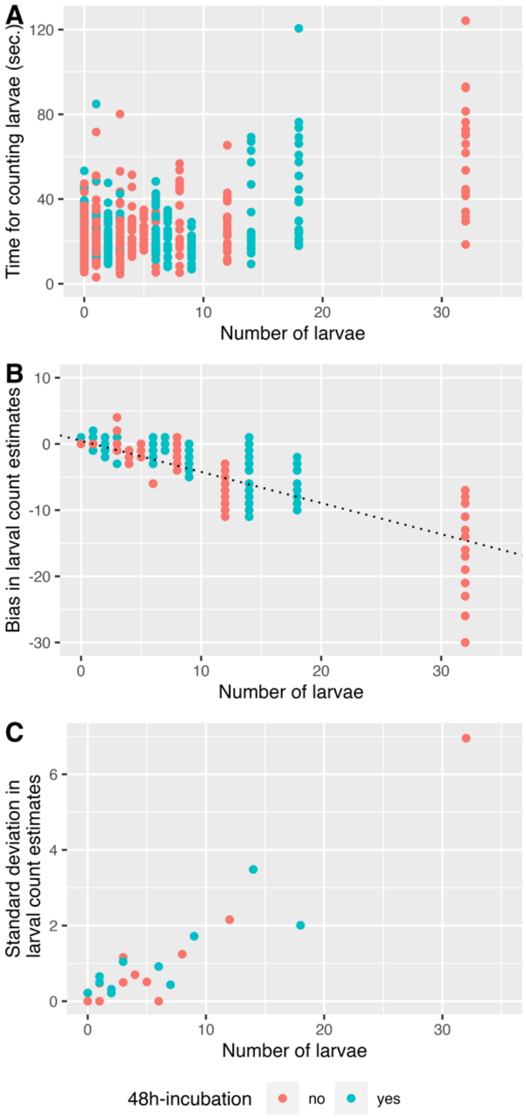


**Fig. S3.** Performance of our new methods in terms of (A) time for counting larvae, (B) accuracy and (C) precision of larval counts in the presence or absence of a 48-hour incubation

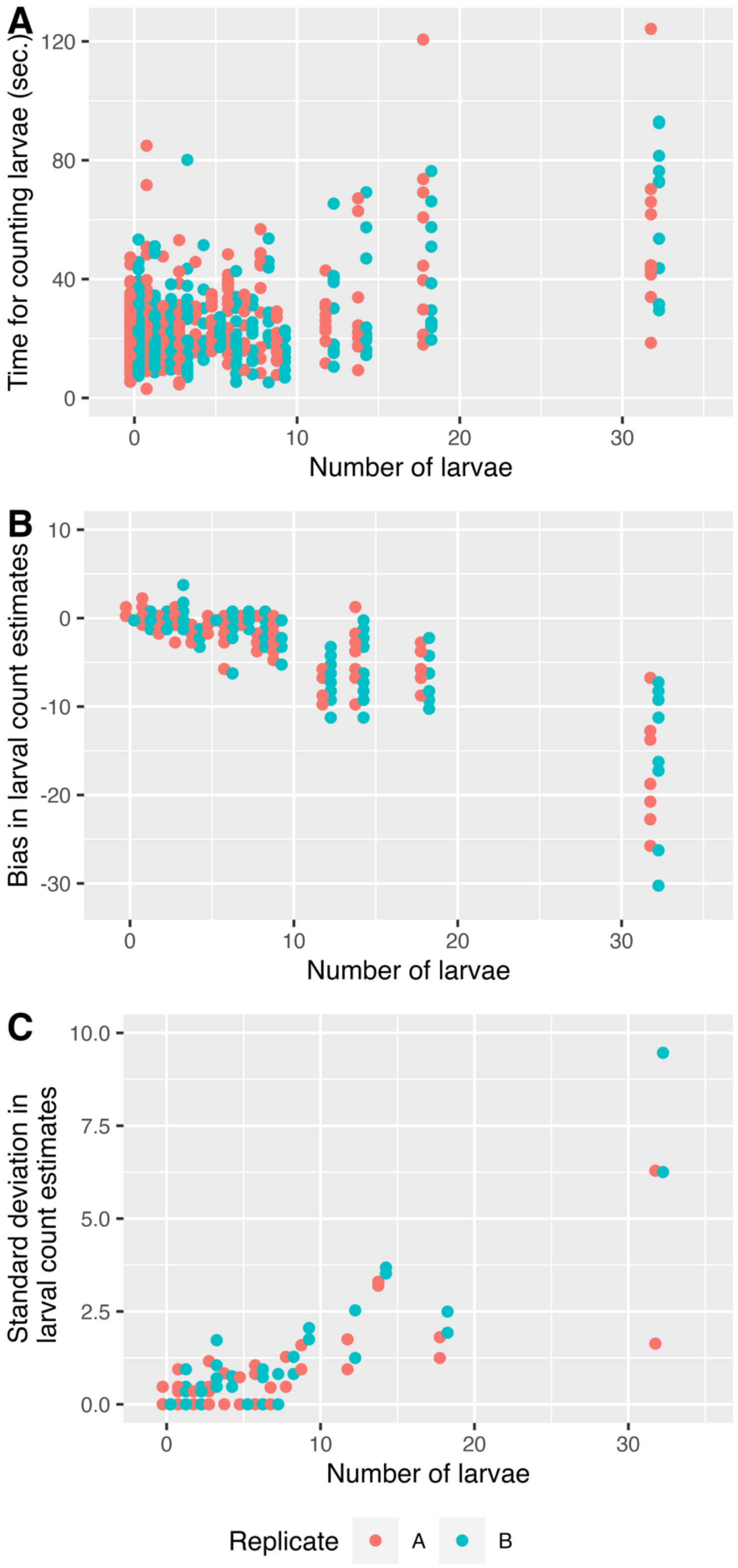


**Fig. S4.** Difference in (A) time for counting larvae but not difference in (B) accuracy and (C) precision of larval counts between the two replicate counts
